# Gut^3^Gel: A High Throughput Mucus Model for Culturing Human Intestinal Microbiota

**DOI:** 10.1101/2025.02.21.639490

**Authors:** Natalia Suárez Vargas, Miguel Antunes, João Sobral, Carolina Silva, Francisco Sousa, Olga Valentina Garbero, Anna Kolková, Livia Visai, Claudio Medana, Sonja Visentin, Paola Petrini, Sebastião van Uden, Daniela Pacheco

## Abstract

The human intestinal microbiota plays a crucial role in health and disease, yet recreating its complex interactions *in vitro* remains a significant challenge. Gut^3^Gel introduced herein as a novel *in vitro* mucus model, designed for culturing complex microbial communities without the need for anaerobic conditions. Intestinal microbiota samples from five donors were individually inoculated in Gut^3^Gel and cultured for 72 hours. Taxonomic composition assessment revealed that Gut^3^Gel sustains diverse microbial species and particularly promotes the growth of mucus-associated bacteria including *Bifidobacterium*, *Lactobacillus*, and *Faecalibacterium*. Microbial metabolic activity within Gut^3^Gel was confirmed by the increased production of acetate and butyrate, as well as of exopolysaccharides. Gut^3^Gel reproduces physiological features of intestinal mucus, providing a reproducible and scalable culturing platform. These features make Gut^3^Gel a promising tool for advancing microbiota research with potential applications in drug screening, microbiome mining, and high throughput testing of microbiome-modulating molecules.

---

The human intestinal microbiota is a dynamic and complex community of microorganisms that plays an integral role in various aspects of health^1^. The microbial distribution along the large intestine is governed by niche-specific factors, such as pH and oxygen gradients, within the colonic mucus layer^2–4^. The mucus is a bilayered structure that covers the colonic epithelial surface: the inner layer, close to the epithelium, is characteristically more viscoelastic and mostly devoid of microbial species, while the outer loose layer, toward the lumen, is less viscoelastic and provides a niche for microbial communities^5,6^. Together, both layers establish complex bidirectional host-microbe interactions and provide protection against pathogenic microbes and harmful luminal compounds^3^. The mucus-associated microbiota composition has a crucial contribution to the regulation of the intestinal mucus barrier function, as microbes influence the properties of the mucus^7^. Therefore, the integrity of the intestinal mucus layer is fundamental for health, as its disruption is linked to the onset of many diseases^7–9^.

Understanding the relationship between microbiota composition and function imbalances, as well as specific diseases remains a complex challenge^10^. Establishing causal links between microbiota and health outcomes is complex due to the dynamic and multi-factorial nature of these interactions. The current gold standard for studying microbiota, *in vivo* animal models, offers valuable insights in establishing causal associations, but faces critical limitations^10^. Ethical concerns, as well as logistical, time, and cost constraints, make widespread use of animal models impractical. Additionally, these models often fail to replicate the human microbiota accurately, as substantial differences exist in the physiology, chemical susceptibility, and microbial colonization between humans and animal models^11^. Another level of complexity lies in the fact that a large portion of the intestinal microbiota is composed of fastidious or yet unculturable microorganisms^12^. A recent evaluation of metagenomic studies revealed that more than 70% of the species cataloged in the human intestinal microbiome lack a cultured representative^13^. This emphasizes the current need for culturing platforms to study microbiota composition, function and interaction in a controlled, sustainable, and reproducible manner.

*In vitro* models have emerged as valuable complementary tools to *in vivo* animal models for studying microbiota^14^. This shift is driven by the growing need to reduce the dependency on animal testing allied to the transition to more efficient screening methods. Widely used systems, with different degrees of complexity, include single or multi-component bioreactors (e.g., TIM-2, SHIME, SIFR, simgi)^15–18^ and microfluidic devices (e.g., intestine-on-chip)^19^. These models have provided valuable insights into the interactions between microbiota, dietary compounds, and therapeutic agents. However, they present some limitations. Bioreactors, while capable of supporting complex microbial communities, are often expensive, labor-intensive, and low throughput^20,21^. Similarly, microfluidic devices excel in simulating fluid dynamics, but are often costly, complex to operate, and limited in scalability^22,23^. These limitations highlight the need for a simpler, scalable, and physiologically relevant platform that balances complexity and throughput, bridging the gap in existing *in vitro* models. In this study, Gut^3^Gel is introduced and studied as a novel versatile *in vitro* model to culture human intestinal microbiota. Gut^3^Gel was designed by reverse engineering key features of the intestinal mucus known to affect microbial dynamics, including chemistry, viscoelastic properties, bilayered structure and gradient of oxygen tension^3^, offering a supporting 3D matrix for microbial adhesion, growth and interaction. The ability of Gut^3^Gel to successfully sustain diverse and stable microbial communities while providing reproducible results across experimental replicates was determined by 16S sequencing, rheological characterisation and gas chromatography–mass spectrometry (GC-MS). To validate the capacity of Gut^3^Gel to maintain a stable microbial community, the composition and metabolic activity of the intestinal microbiota from five healthy donors were evaluated. As a comparison, Brain Heart Infusion (BHI), a widely used non-selective medium for culturing intestinal microbiota, including fastidious microorganisms, was used as a control^12,24^. The findings presented here establish Gut^3^Gel as a promising *in vitro* model for advancing microbiota research and its application in health and disease.

## Results

### Gut^3^Gel: A Mucus Biomimetic Gradient Hydrogel for Intestinal Microbiota Cultivation

Gut^3^Gel was designed to reflect the multitude of microenvironments exhibited by the human intestinal mucus. Gut^3^Gel displays a gradient structure with a stiffer, densely crosslinked region stained in pink, and a softer, less crosslinked region stained in yellow (Fig. 1a), reflecting the attached and loose layers of native colonic mucus, respectively. The staining was used to depict the heterogeneous structure of Gut^3^Gel, which specifically binds the crosslinking ions. This gradient in crosslinking density is linked to a progressive decrease in oxygen tension, starting from air oxygen tension (∼220 µmol.L-1) at the Gut^3^Gel surface and reaching a minimum oxygen tension of 120 µmol.L-1 at deeper layers (Fig. 1b). The physicochemical properties of Gut^3^Gel were also assessed through rheological testing (Fig. 1c). The gel-*like* structure was validated by an average conservative modulus (G’) of about 500 Pa and a dissipative modulus (G’’) of about 80 Pa across the full range of tested frequencies. These features sustained the growth in Gut^3^Gel of approximately 83% of the bacterial genera identified in the original samples of the donated intestinal microbiota (Fig. 1e).

**Fig. 1.**
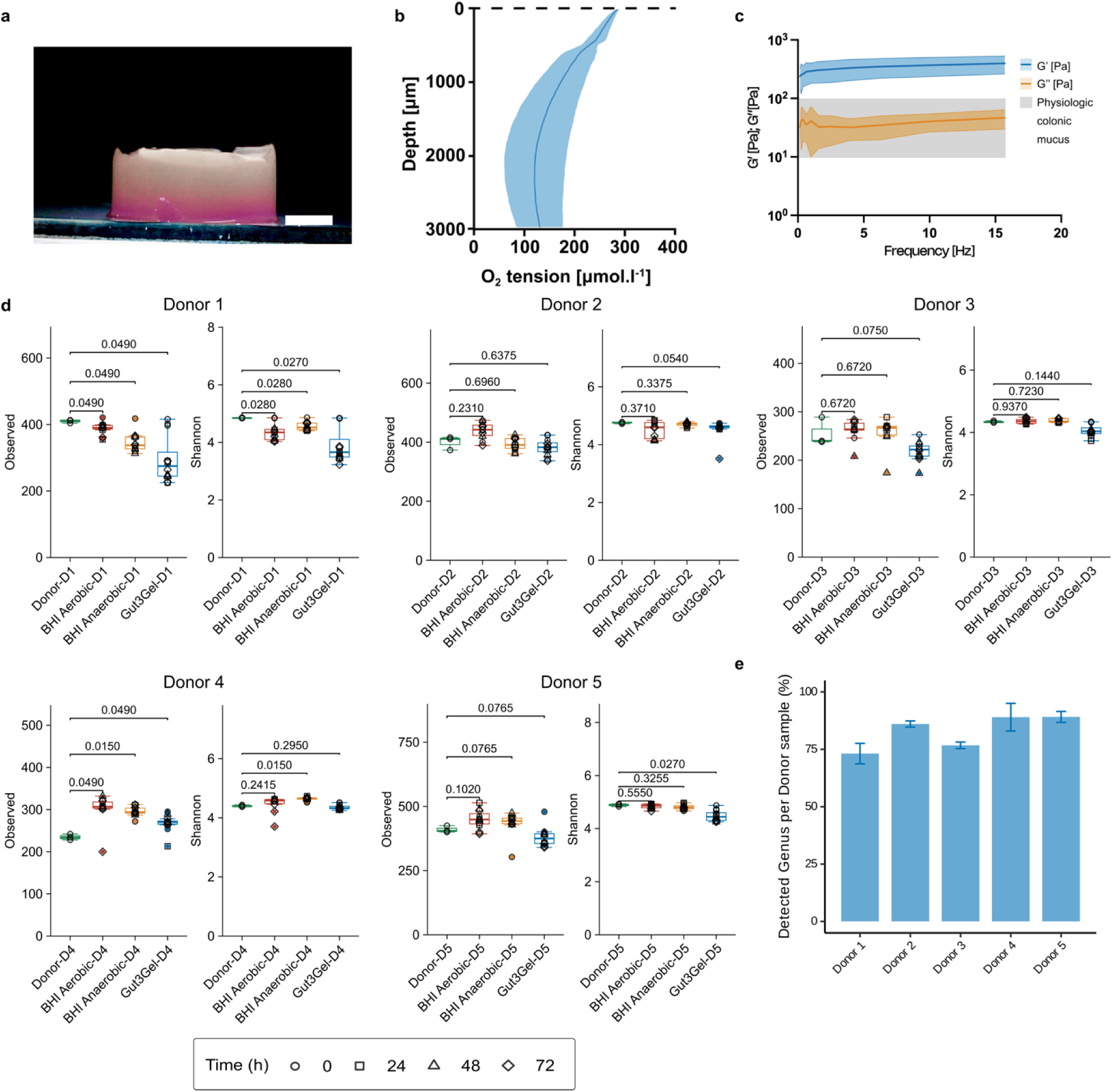
Gut3Gel characteristics and performance to sustain *in vitro* intestinal microbiota. **a-c,** Topographical and physical features of Gut^3^Gel. **a,** Macroscopic image of Gut^3^Gel stained with alizarin red. The scale bar corresponds to 3 mm. **b,** Viscoelastic properties of Gut^3^Gel: conservative, G□ (Pa; blue line), and dissipative moduli, G□□ (Pa; orange line) of Gut^3^Gel (n ≥ 6). **c,** Heterogeneous distribution of O2 tension through the structure of sterile Gut^3^Gel (n = 5). The shaded ranges in **b, c,** correspond to the standard deviation. **d,** Alpha diversity analysis of microbial communities in Gut^3^Gel. Alpha diversity was determined based on observed ASVs and the Shannon index to evaluate microbial diversity and richness of the microbial communities within Gut^3^Gel and BHI (under aerobic or anaerobic conditions) cultured for 24, 48, or 72 h, in comparison to the inoculum (donor samples). The Benjamini–Hochberg adjusted p-values are shown between culturing platforms and the donor samples (Wilcoxon rank-sum test). **e,** Percentage of genera identified in Gut^3^Gel after 24 h of culture of microbiota from five different individuals.

### Gut^3^Gel Supports the Growth of Diverse Microbial Communities

The evolution of microbial communities was monitored using 16S rRNA sequencing of samples collected from Gut^3^Gel at four time points (0, 24, 48, and 72 h) and the initial fecal inocula (donor samples). Microbial diversity and richness within samples were assessed by alpha diversity measures. Based on observed Amplicon Sequence Variants (ASVs) in Gut^3^Gel, the diversity was comparable to the starting inoculum for all donors, except Donors 1 and 4, suggesting a general conservation of ASVs during culturing in Gut^3^Gel (Fig. 1d). Richness and evenness were further evaluated using the Shannon diversity index. Shannon index values remained relatively stable for Donors 2 and 4, while slightly decreased for Donor 3. Donors 1 and 5 displayed the most substantial reduction in diversity, reflecting a shift in the microbial community between the initial timepoint (0 h) and subsequent timepoints (24, 48 and 72 h). This decrease in diversity indicates that culturing in Gut^3^Gel is either driving the loss of bacterial species or leading to significant alterations in the abundances of specific taxa when compared to the initial inoculum.

To understand the factors driving the observed differences in diversity, the microbial composition of donor samples was compared to that of samples incubated in the BHI medium under aerobic and anaerobic conditions. At the phylum level, microbial communities in Gut^3^Gel were predominantly composed of Firmicutes (54–74%), followed by Actinobacteriota (9–32%), Bacteroidota (8–29%), Proteobacteria (0.4–3.4%), and Verrucomicrobia (0–2.4%) (Fig. 2). The fecal inocula from the five donors varied in their Firmicutes-to-Bacteroidota ratios, which were 3.3, 4.1, 2.5, 2.2 and 4.2 for Donors 1, 2, 3, 4, and 5, respectively. Culturing in Gut^3^Gel led to an average increase in Firmicutes abundance of 1.2-fold, 1.5-fold, and 1.7-fold in this ratio across donors after 24, 48, and 72 h, respectively. Besides the increase in Firmicutes abundance, the most notable alteration was the substantial increase in Actinobacteriota abundance, which was 2- to 10-fold higher in Gut^3^Gel compared to the faecal inocula. Compared to the BHI medium, the microbial communities in Gut^3^Gel displayed lower abundance of Proteobacteria. In BHI, Proteobacteria were more abundant under aerobic conditions (7–29%) than under anaerobic conditions (2–10%), suggesting that the presence of oxygen is promoting the growth of facultative anaerobes.

**Fig. 2.**
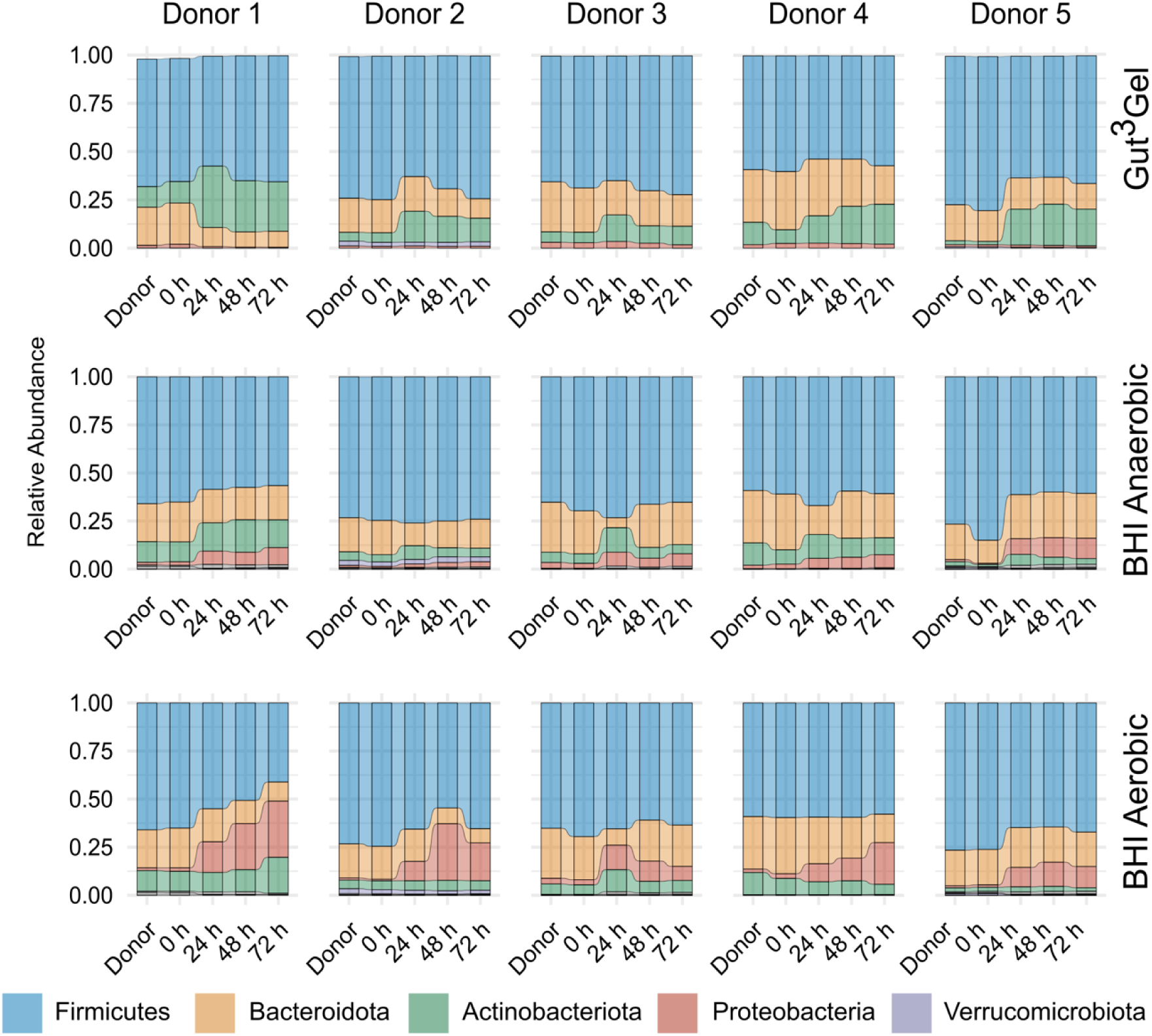
Taxonomic composition of intestinal microbiota cultured in Gut^3^Gel. Barplots display the relative abundance at the phylum level from the intestinal microbiota cultured in Gut^3^Gel for up to 72 h compared to the BHI medium (under aerobic and anaerobic conditions).

Besides microbial composition, it is relevant to investigate how much the microbial communities deviate from the inocula and between culturing platforms. Beta diversity analysis shows a significant departure of Gut^3^Gel from the planktonic condition (BHI medium) in terms of dissimilarity both based on abundance and phylogeny (weighted UniFrac) across all donors (PERMANOVA, *p* < 0.05) (Fig. 3a-e; Supplementary Table S1). Principal coordinate analysis (PCoA) of microbial profiles in Gut^3^Gel showed clustering of samples by time points with minimal divergence across all donors (Fig. 3a-e). This is supported by the absence of significant alterations over time of the dissimilarity measure weighted UniFrac (PERMANOVA, *p* > 0.05) between replicates across timepoints (24 to 72 h) in Gut^3^Gel. Beta diversity was also assessed among donor samples, with Donor 1 displaying the highest dissimilarity relative to the other donors (Fig. 3f).

**Fig. 3.**
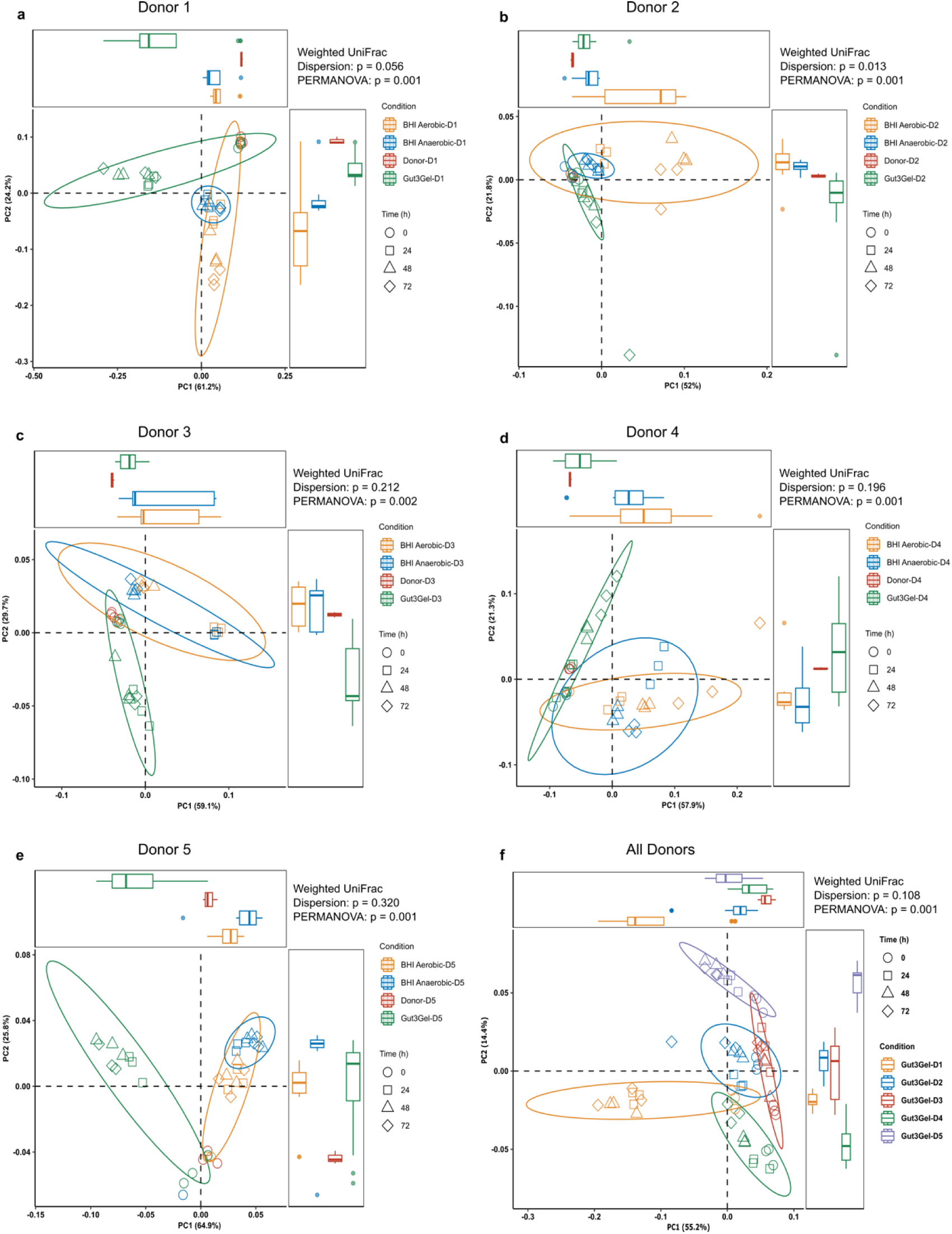
Beta diversity analysis. **a - e,** Beta diversity was determined based on the dissimilarity metric Weighted Unifrac to evaluat the dissimilarities in taxonomic composition between the microbial communities in Gut^3^Gel and BHI (under aerobic and anaerobi conditions) for donors 1, 2, 3, 4, and 5, **a** to **e**, respectively. **f,** Dissimilarity analysis of taxonomic compositions of intestinal microbiota from the five individual donors cultured in Gut^3^Gel. Significant differences between microbial communities acros samples were tested using PERMANOVA. Adjusted p-values and β-dispersion values are displayed for each comparison. Ellipses represent 95% confidence intervals. Side boxplots display the distribution of the samples along each principal coordinate axis, providing an overview of culturing condition dispersions.

The microbial composition at the Family taxonomic level in Gut^3^Gel was compared across all donors to identify which families are promoted independently of the donor sample (Fig. 4a). Species belonging to the Families *Bacteroidaceae*, *Coriobacteriaceae*, *Prevotellaceae*, *Lachnospiraceae*, *Oscillospiraceae*, and *Ruminococcaceae* (the latter 3 being mucosal-associated) consistently exhibited a substantially high abundance across all donors, with a centered log-ratio (CLR) value above 2, indicating their abundance is more than seven times the geometric mean. The high variability in CLR abundance of species from several Families, as evidenced by the distinct microbial abundance profiles across donors, underscores the ability of Gut^3^Gel to preserve interindividual differences among donors. Moreover, compared to BHI anaerobic, Gut^3^Gel seems to further promote and sustain the growth of the dominant species from the donor samples (Supplementary Fig. S1).

**Fig. 4.**
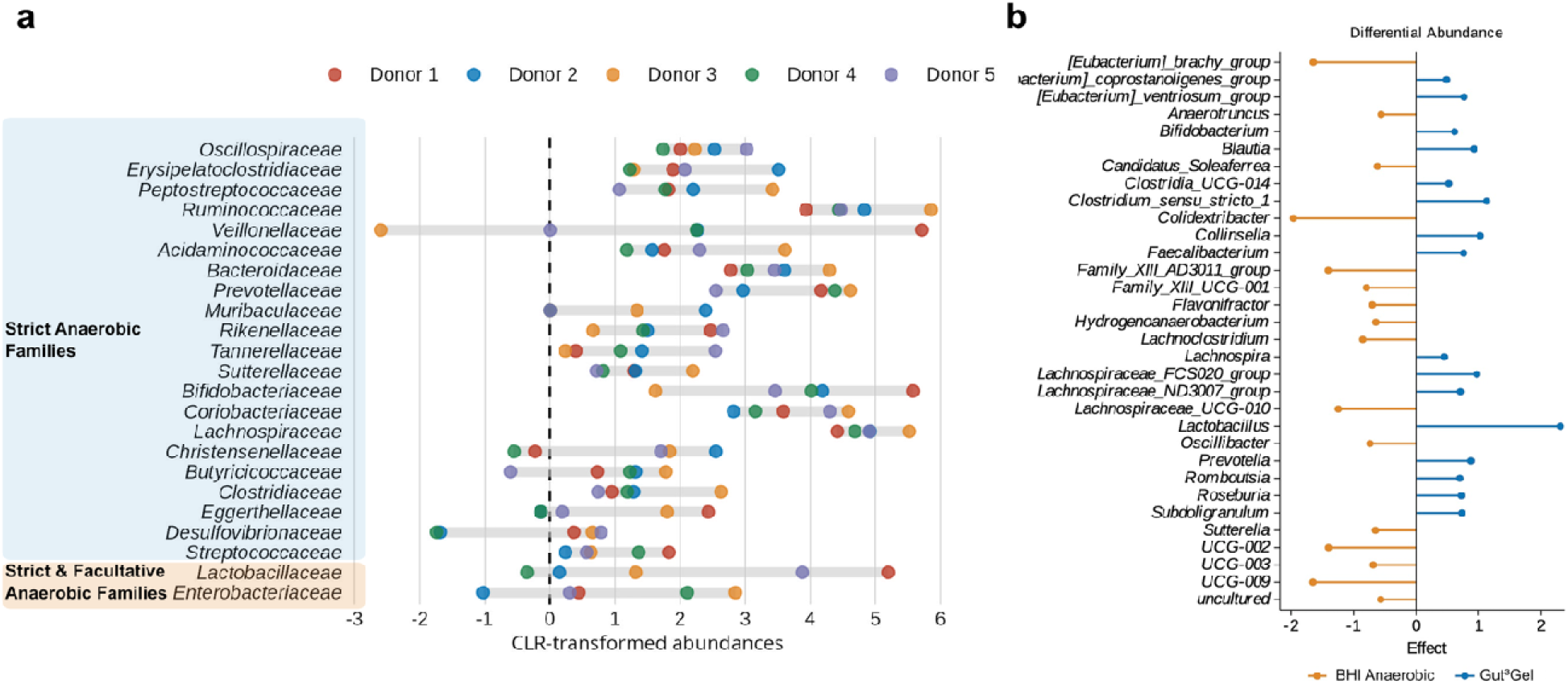
Comparison of microbial composition among donor samples. **a,** Comparison of the microbial clr-transforme abundances at the Family taxonomic rank in Gut^3^Gel among donor samples. **b,** Differential abundance analysis using ALDEx between Gut^3^Gel and BHI (anaerobic) across all donor samples and culturing times ranging from 24 to 72 h.

Differential abundance analysis was conducted to identify bacteria that significantly differ based on pairwise comparisons of microbial relative abundances in Gut^3^Gel and BHI anaerobic culturing platforms across all donors between 24 and 72 h of culturing. Bacterial species belonging to genera *Lactobacillus*, *Clostridium sensu stricto*, *Collinsella*, *Prevotella*, *Bifidobacterium*, *Roseburia* and *Blautia* were highly abundant in Gut^3^Gel compared to BHI (Fig. 4b). Bacteria from these genera are associated with mucin and polysaccharide metabolization and short-chain fatty acid (SCFA) production. In contrast, BHI exhibited higher abundances of genera such as *Eubacterium brachy group*, *Colidextribacter*, *Lachnoclostridium*, *Oscilliba*cter, and *Sutterella* (Fig. 4b). These differences underscore the influence of the culturing platform on microbial community composition, with Gut^3^Gel, characterized by its mucus-mimicking properties, favoring the growth of mucus-associated microbes. On the other hand, the nutrient-rich, planktonic environment of BHI supports bacterial communities adapted to more general nutrient conditions.

### Alterations in Gut^3^Gel structure and SCFA production underscore dynamic microbial interactions

The metabolic activity of intestinal microbiota can be assessed by monitoring changes in the structural and rheological properties of the Gut^3^Gel matrix (Fig. 5a). As bacteria grow, metabolization of Gut^3^Gel results in the production and secretion of viscoelastic compounds, particularly exopolysaccharides (EPS), which is supported by the observed increase in viscoelastic moduli through rheological assessment. The dynamic interactions between microbial activity and physical properties of the environment are evidenced by the increase in abundance over time of EPS-producing bacteria in Gut^3^Gel (Fig. 5b), and the increase of both conservative (G’) and dissipative (G’’) moduli of Gut^3^Gel over the 72 h of culturing, except for Donors 2 and 3 that peak at 24 h (Fig. 5a). Curiously, further corroborating the proposed hypothesis, when analyzing Donor 3 samples, *Lactobacillaceae* were not detected and *Bifidobacteriaceae* were present in very low abundance, which was concomitant with the absence of an increase of viscoelastic properties of Gut^3^Gel after 24 h of culturing. Additionally, the sharp increase in the viscoelastic properties of Gut^3^Gel in Donor 4 at 72 h of culturing may be associated with the substantial increase in *Lactobacillus* abundance between 48 and 72 h (Fig. 5b).

**Fig. 5.**
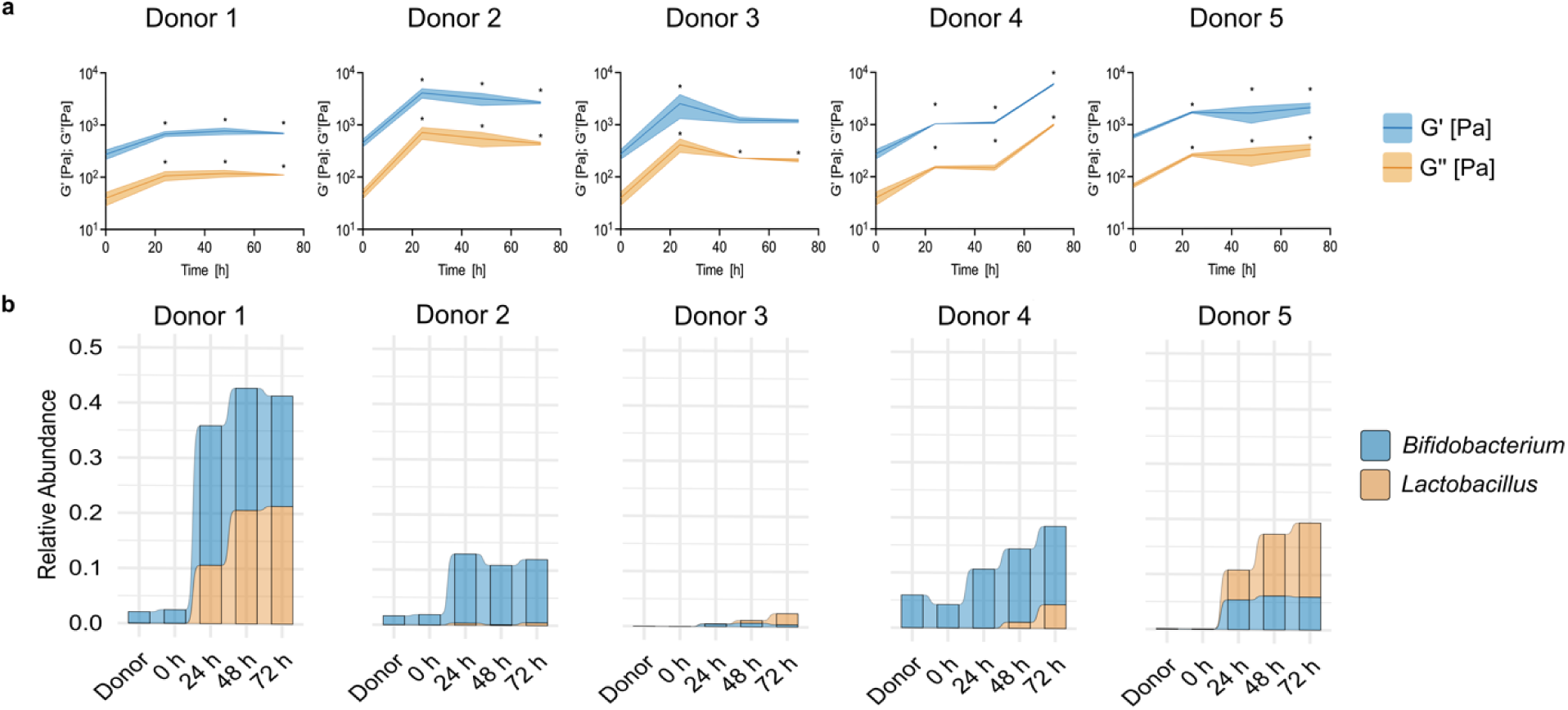
Assessment of physical properties of Gut^3^Gel in the context of microbial activity dynamics. **a,** Rheological characterization, including conservative, G□ (Pa; blue line), and dissipative moduli, G□□ (Pa; orange line) of Gut^3^Gel inoculate with intestinal microbiota from five different donors. The shaded range corresponds to the standard deviation. The adjusted p-value are shown between time points (ANOVA followed by Bonferroni correction for multiple comparisons). **b,** Relative abundance of *Lactobacillus* and *Bifidobacterium* during the cultivation of intestinal microbiota from five different donors in Gut^3^Gel.

The metabolic activity of intestinal microbiota was also monitored through the quantification of short-chain fatty acids (SCFAs). The production of SCFAs, in particular acetate, propionate and butyrate, was assessed by quantifying their concentrations over time in both Gut^3^Gel and BHI (under anaerobic conditions). Despite the covariance among donor samples being high (above 20%), the results demonstrate that the SCFA production profiles display a consistent trend across all donor samples. Compared to BHI under anaerobic conditions, Gut^3^Gel displayed higher levels of acetate and butyrate, whose production was favored over propionate (Fig. 6). In Gut^3^Gel, the acetate/propionate/butyrate ratios across donors shifted from 2.5:1:3 at 24 h to 2.5:1:3.5 at 72 h, reflecting an increase in the proportion of butyrate, while in BHI the ratio was maintained at 1:1:1 from 24 to 72 h. Acetate levels in Gut^3^Gel increased on average 10-fold (from 38 ± 44 to 411 ± 157 µg.mL-1) in the first 24 h and then either stabilized or slightly decreased over the following 48 h. In contrast, butyrate levels in Gut^3^Gel rose on average 20- fold (from 19 ± 15 to 472 ± 148 µg.mL-1) during the first 24 h and continued increasing steadily throughout the 72 h culturing period. The concentration of produced SCFAs for each individual donor sample can be found in Supplementary Fig. S2. The results from SCFA production, in general, are in agreement with the most abundant species detected in Gut^3^Gel (Supplementary Fig. S3). Donor 1 displayed a high abundance of acetate and butyrate producers, such as *Bifidobacterium* (30%), and *Megasphaera* (30%), respectively. Donor 2 displayed a high abundance of acetate-producers such as *Bifidobacterium* (13%), *Ruminococcus* (13%), and *Prevotella* (7%), and the butyrate producer *Faecalibacterium* (15%). As for Donor 3, the microbial community was mainly composed of the butyrate-producer *Faecalibacterium* (35%) and the acetate-producer, *Prevotella* (11%). The sharp increase in acetate in Donor 4 is concomitant to the high abundance of acetate-producers such as *Bifidobacterium* (14%), *Prevotella* (19%), while for Donor 5 there is a high abundance of *Lactobacillus* (11%) and *Bifidobacterium* (6%). These results highlight the ability of Gut^3^Gel to support distinct SCFA production dynamics coupled with the microbial community composition.

**Fig. 6.**
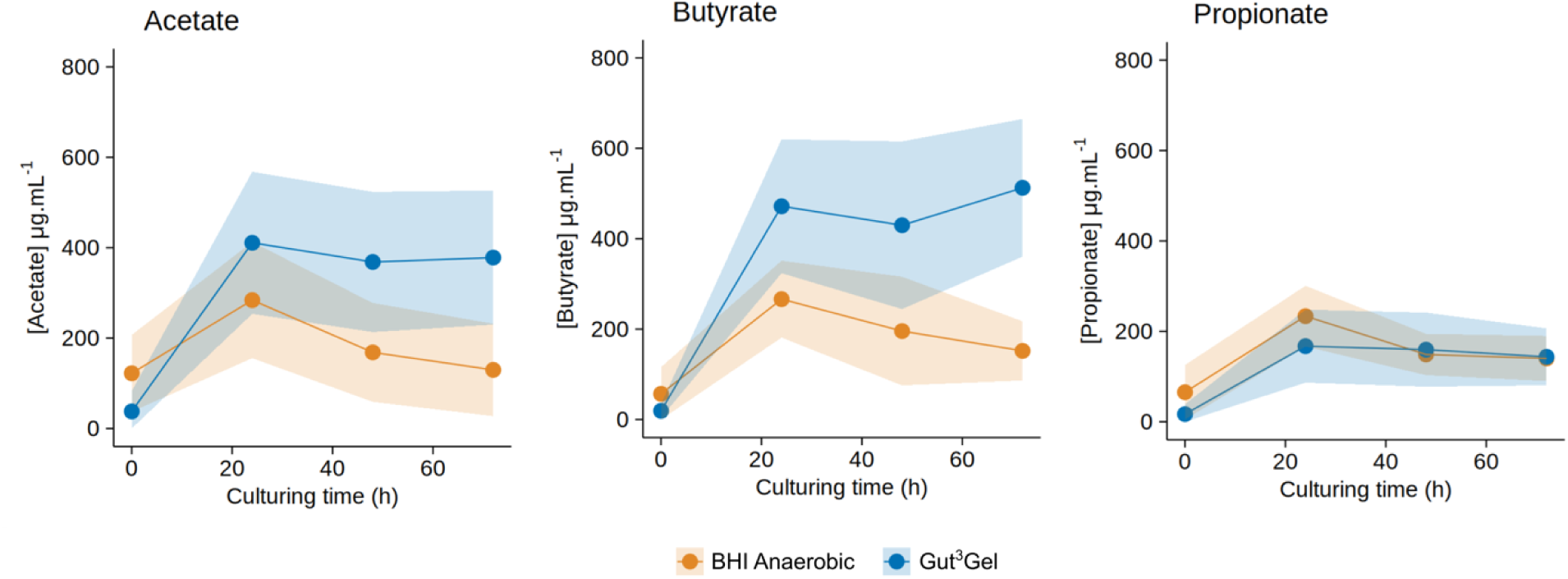
Production of SFCAs in Gut^3^Gel over time. Levels of SCFAs (acetic acid, butyric acid and propionic acid) produced by the intestinal microbiota inoculated in Gut^3^Gel. The data represents the average concentration of each SCFA of all microbiota cultured in either Gut^3^Gel (blue line) or BHI under anaerobic conditions (orange line). The shaded range corresponds to the standard deviation.

## Discussion

This study introduces and validates Gut^3^Gel as a novel *in vitro* 3D model of the human intestinal mucus environment that supports the growth of intestinal microbiota. Using 16S rRNA gene amplicon sequencing and targeted metabolite analysis, it is demonstrated that Gut^3^Gel sustains diverse microbial communities over 72 hours of culturing time without the need for anaerobic conditions.

Gut^3^Gel is a scalable, user-friendly culturing platform designed to replicate the 3D architecture and physiological conditions of intestinal mucus. It mimics key physicochemical properties of colonic mucus, including the presence of 50 mg.mL^-1^ mucin, as observed in physiological conditions^25^, and a mucus architecture comprising a loose layer that hosts microbiota, alongside a highly crosslinked (attached layer) region. Similar to the mucus model we have previously proposed^26^, the gradient structure of Gut^3^Gel establishes a gradual decrease in oxygen tension across the thickness of Gut^3^Gel, creating a biomimetic environment capable of hosting anaerobic bacteria strains. The flexible chemical composition of Gut^3^Gel can be exploited to include specific culture medium, as previously described^26^. In this work, Gut^3^Gel was produced within BHI, which, together with its intrinsic physicochemical characteristics, creates a microenvironment conducive to microbiome growth.

The viscoelastic properties of colonic mucus typically range from 10-100 Pa for the conservative component (G’), with characteristic tan δ values below 1^27–29^, depending on hydration, diet, microbiota composition, and health status. Gut^3^Gel was designed with a G’ greater than 100 Pa and a tan δ of 0.2, corresponding to a dissipative modulus of 20 Pa, ensuring substrate stability that sustains microbial growth for over 72 h. These characteristics enabled the growth of approximately 83% of the genera found in the original donor samples, including strictly anaerobic bacteria, while promoting the proliferation of mucus-adherent microbiota.

One of the most striking differences between the microbial composition of Gut^3^Gel and BHI is the substantially lower abundance in Gut^3^Gel of Proteobacteria, mainly *Escherichia* and *Pseudomonas* and notably higher levels of Actinobacteriota, with *Bifidobacterium* as dominant species. In *in vitro* models, substantial increases in the abundance of *Bacteroides* and *Enterobacteriaceae* genera are commonly observed^30,31^. This phenomenon is often partly attributed to challenges in establishing effective oxygen gradients, a recurring challenge in accurately recreating *in vitro* intestinal microbiota models. The impact of oxygen stress *in vitro* can be inferred by evaluating the growth patterns of facultative anaerobes, such as enterobacteria or lactobacilli, as well as extremely oxygen-sensitive (EOS) species^32^. In BHI under aerobic conditions, the effect of oxygen is evident, especially for Donors 1 and 2, where enhanced growth of *Escherichia* and *Pseudomonas* is observed. Under anaerobic conditions, *Escherichia* is also detected in BHI, although at lower abundances compared to aerobic conditions, suggesting some degree of oxygen stress. In contrast, Gut^3^Gel exhibits a distinct microbial profile. Mucus-associated bacteria, such as lactobacilli^33^, were detected in almost all donors, but in higher abundances in Donors 1 and 5, while *Escherichia* and *Pseudomonas* were absent, except for Donors 3 and 4. The presence of *Escherichia* was detected in low abundances in Donors 3 and 4, which may not necessarily indicate oxygen stress but rather result from the formation of a microaerophilic environment created by the oxygen gradient within Gut^3^Gel. Another hypothesis is the reduced antimicrobial activity by species like *bifidobacteria* and *lactobacilli*^34^, which were present in high abundance in all donors, but almost absent in Donor 3. Nevertheless, the reduced presence of facultative anaerobes, allied to the proliferation of EOS species in Gut^3^Gel, underscores its capacity to sustain the growth of anaerobes from the intestinal microbiota. Remarkably, this was achieved without the need for strict anaerobic incubation, highlighting the potential of Gut^3^Gel as an effective and practical *in vitro* model for studying and monitoring intestinal microbiota.

Metabolic activity in Gut^3^Gel was evidenced by alterations in its viscoelastic properties during the culture period, as well as the production of SCFAs. In Gut^3^Gel, the concomitant and substantial increase in acetate and butyrate was in agreement with the high abundance of the acetate producers, such as *Bifidobacterium* species, and butyrate-producers such as *Faecalibacterium*. The increased abundance of *Faecalibacterium* in Gut^3^Gel, despite the absence of anaerobic conditions, is noteworthy. While these bacteria are highly prevalent in the human intestine, their isolation and culture remain challenging due to their extreme oxygen sensitive (EOS) nature^35^. The production of butyrate by *Faecalibacterium* and other butyrate-producers is driven by acetate consumption through cross-feeding interactions with acetate-producing bacteria such as *Bifidobacterium*^35,36^. In fact, the generally lower levels of acetate in Donor 3 might be attributable in part to the very low abundance of *Bifidobacterium* compared to the other Donors. Furthermore, Donor 1 showed the highest levels of butyrate, which might be explained by cross-feeding interactions taking place between the acetate and lactate producers, such as *Bifidobacterium* and *Lactobacillaceae* and butyrate producers, particularly *Megasphaera*^37^.

The substantial modifications of Gut^3^Gel matrix may be attributed to mucin metabolization and exopolysaccharide production. These dynamic interactions reflect the functional interplay occurring *in vivo*, where microbial activity is continually reshaping the mucosal environment. The mucin-based composition of Gut^3^Gel promoted the growth of physiologically mucus-associated bacteria, including *Bifidobacteriaceae*, *Lactobacillaceae*, *Lachnospiraceae*, *Ruminococcaceae* and *Faecalibacterium*^38–40^. This microbial selection closely aligns with *in vivo* data, where mucus-associated microbiota is dominated by families like *Bifidobacteriaceae*, *Lactobacillaceae*, *Ruminococcaceae* and *Lachnospiraceae*, while families like *Bacteroidaceae* and *Enterobacteriaceae* are more prominent in the lumen^41^. Similar trends have been observed in established *in vitro* models that incorporate mucin such as M-SHIME, which shows a high prevalence of Firmicutes and Actinobacteriota and increased butyrate production upon mucin supplementation^42,43^. The presence of mucin in Gut^3^Gel not only supports the growth of *Lactobacillaceae* and *Bifidobacteriaceae*, but also suggests that the establishment of specific bacterial populations may rely on the distinct microenvironments created within the *in vitro* platform for optimal growth.

In summary, the Gut^3^Gel mucus-mimicking 3D structure represents a versatile and scalable tool that advances microbiota research by addressing critical challenges in replicating the intestinal mucus environment for culturing complex microbial communities, including mucus-associated microbes. Its high-throughput compatibility and ability to function without anaerobic conditions enable the simultaneous testing of multiple conditions using a large number of donor samples with various replicates. This is crucial for preserving interindividual differences, as it eliminates the need to pool samples. Additionally, the experimental parameters of Gut^3^Gel can be finely tuned to specific research objectives by incorporating growth media, specific nutrients, and growth factors directly into the Gut^3^Gel structure. The findings presented in this study provide a foundation for exploring the impact of therapeutic molecules on complex microbial community interactions within a physiologically relevant *in vitro* model. Gut^3^Gel shows considerable potential for future applications, including drug screening, microbiome mining, permeability assays, and other biomedical research areas involving host-microbiome interactions.

## Methods

### Donor Recruitment

All donors provided written informed consent, confirming their understanding of the study and their voluntary participation. Approval from an ethical committee was not required for this study, as it did not involve prospective evaluation or the use of laboratory animals. The study was conducted exclusively employing non-invasive methods, specifically the collection of faecal samples. Furthermore, all donors’ personal information was kept anonymous upon sample collection and throughout data analysis.

### Gut^3^Gel

Gut^3^Gel gradient colonic (G3GG colonic, Bac3Gel Lda) was adopted as it captures key features of the gut environment, such as chemical composition, gradient structure, and viscoelastic properties. Gut^3^Gel is a hydrogel-based structure composed of 50 mg.mL-1 porcine stomach type III mucin (Sigma-Aldrich, M1778; Germany) that falls within the ranges previously determined in intestinal mucus^25^ and it was produced in Brain Heart Infusion medium (BHI, Sigma-Aldrich). Gut^3^Gel colonic exhibits a gradient structure that is tightly connected with a 3D gradient of oxygen tension. The method of production is patented (IT102018000020242A)^44^.

### Isolation and culturing of intestinal microbiota

Intestinal microbiota was isolated from faecal samples from five healthy individuals following the European Guidelines of Faecal Matter Transplantation^45^. Briefly, a minimum amount of 30 g of faecal sample was suspended in 150 mL of 0.9% NaCl using a blender, followed by a filtration step. All collections were performed in the morning and immediately used upon isolation (maximum 30 minutes after processing). All donors were European, aged between 25 to 36 years, with a BMI ranging from 18.5 to 30, without gastrointestinal complaints or diagnosed disorders, and free of antibiotics or food supplements at least 3 months prior to this study. Isolates of intestinal microbiota were cultured in Gut^3^Gel for 24, 48, and 72 h. Cultures in Gut^3^Gel were prepared by inoculating the intestinal microbiota on top of Gut^3^Gel in a 1:1 ratio, followed by incubation at 37 ◦C under aerobic conditions (no anaerobic conditions were applied during the incubation period when Gut^3^Gel was exploited).

For comparison, planktonic cultures were conducted in parallel by inoculating the microbiota into BHI medium, while maintaining the same volume proportions. These planktonic cultures were incubated under either aerobic or anaerobic conditions for 24, 48, and 72 h, and served as experimental controls. Anaerobic conditions in the BHI cultures were established using AnaeroGen Gas packs (Oxoid AN0025A), a method shown to provide conditions comparable to an anaerobic chamber for cultivating intestinal microbiota^46^.

Sterile Gut^3^Gel and BHI were included as additional controls of sterility, rheological properties, oxygen tension, and sequencing.

### Rheological characterization

To determine the effect of intestinal microbiota colonization and metabolization, the viscoelastic properties of Gut^3^Gel colonized with intestinal microbiota for different culture periods (0, 24, 48, and 72 h) were evaluated using an Anton Paar MCR501 Rheometer (Austria) with a 25 mm diameter parallel plate geometry (serial number 79044/7810) at 25 ◦C. The linear viscoelastic region (LVR) was determined through strain sweep analyses employing a strain logarithmic ramp varying from 0.1% to 1000% at a frequency of 1 Hz. Oscillatory frequency sweeps were further performed to evaluate both conservative, G’, and dissipative, G’’, moduli at 0.5% (at strain amplitudes within the linear regime) with frequencies changing logarithmically in the 0.1-20 Hz range. Statistical significance was assessed using ANOVA with a significance level of 0.05 in GraphPad Prism version 10 (GraphPad Software, USA). The Bonferroni correction was applied as a post-hoc test for multiple comparisons.

### Microbial community composition analysis

For each time point, the supernatant of Gut^3^Gel was discarded and Gut^3^Gel was washed only to consider the bacteria that colonized its structure. To retrieve the bacteria within Gut^3^Gel, 50 mM of sodium citrate (Sigma) was added to Gut^3^Gel (dilution factor = 2.5) to fully dissolve its structure. The resulting solution was transferred into 2 mL microtubes, followed by 10 minutes of centrifugation at 14,000 rpm. The supernatant was used for SCFA quantification and the resulting pellet underwent DNA extraction. DNA extraction of all samples (Donor, dissolved Gut^3^Gel, BHI aerobic and anaerobic) was performed following the QIAamp Power Fecal Pro DNA Kit guide (QIAGEN) with minor adaptations on the pre-processing step. The pellet was resuspended with 800 µL of the CD1 solution from the QIAamp Power Fecal Pro DNA Kit, transferred to the PowerBead Pro Tube, and proceeded accordingly to the manufacturer.

To assess microbial community composition, the V3-V4 hypervariable regions of the 16S ribosomal RNA (rRNA) gene were targeted for bacterial identification, using primer pairs referenced by Klindworth et al. and Herlemann *et al.*^47,48^, and following the 16S Metagenomic Sequencing Library Preparation Illumina protocol (Part # 15044223 Rev. B, Illumina, CA, USA). The ZymoBIOMICS Microbial Community Standard (Catalog D6306, Zymo Research, Irvine, CA, USA) was utilized as a quality control assessment to evaluate potential biases and artifacts in the PCR and sequencing steps. Library preparation and amplicon sequencing (Illumina Miseq, 2x300 cycles) were performed in the NGS Lab at Bac3Gel, Portugal.

The raw paired-end reads were primer-trimmed using cutadapt. Analysis of the 16S amplicon sequences was performed using QIIME2 (version 2024.5) and R (v4.4.2) in RStudio (version 2024.09.1). Denoising and generation of amplicon sequence variants (ASVs) was performed using the DADA2 plugin within QIIME2. Taxonomic assignment of ASVs were determined with the q2 feature-classifier plugin against the SILVA database version 138.2 using a classifier trained on the V3-V4 regions. Low-abundance ASVs that were present with a total ASV count of less than 0.1% the maximum count were excluded. Furthermore, ASVs that were taxonomically unclassified at phylum rank or taxonomically assigned to mitochondria or chloroplast were removed from the abundance table.

Alpha diversity analysis (Shannon diversity index) was performed using the R package phyloseq v1.46.0. Beta diversity analysis and ordination plots were created using the R packages phyloseq and ape v5.8. Principal coordinate analysis (PCoA) was employed as the ordination method, with weighted UniFrac used as the distance measure. For compositional data analysis, zero values in the abundance table were replaced using the count zero multiplicative method with the R package zCompositions v1.5.0.3. The data was then transformed using the centered log-ratio (CLR) method. Exploratory data analysis was conducted through compositional principal component analysis (PCA) of the datasets, achieved via singular value decomposition (SVD) of the CLR-transformed data.

### Quantification of Short-Chain Fatty Acids (SCFA)

The quantification of SCFAs produced was performed in parallel with DNA extraction starting from the solution resulting from the dissociation with 50 mM of sodium citrate of Gut^3^Gel cultured with microbiota. This solution was centrifuged (10 minutes at 14,000 rpm), followed by the lyophilization of 50 µL of each supernatant sample, which was further processed for quantification. Briefly, for derivatization, 500 µL of BSTFA (N,O-Bis(trimethylsilyl)trifluoroacetamide) [100 µg.mL-1] prepared in dichloromethane (DCM) were added to each sample and the lyophilized samples were incubated at 52◦C for 40 minutes. Following centrifugation (5 minutes, 3500 rpm, room temperature), the control samples were directly analyzed by Gas chromatography–mass spectrometry (GC-MS), while the other samples were diluted in DCM^49^. The concentration of the SCFAs content in each sample was determined using a Nexis GC-2030 equipped with an autoinjector and a capillary column and detected on a single quadrupole mass spectrometer with an electron ionization source. Quantification was performed through the construction of a calibration curve specific for each compound.

### Statistical Analysis

R was used for statistical analysis unless stated otherwise in the Methods section. Details of the applied statistical tests are provided in the figure legends. Evaluation of significant differences in species richness and Shannon diversity between culturing platforms and the donor samples was performed using Wilcoxon rank-sum tests. To evaluate significant differences between microbial communities across samples, permutation analysis of variance (PERMANOVA) was performed. Beta dispersion between groups was tested using the vegan v2.6.6.1 package, as PERMANOVA assumes homogeneity of dispersions. Pairwise PERMANOVA using the RVAideMemoire v0.9.83.7, was used for pairwise comparisons between culturing platforms and culturing times. The p-values were adjusted using the Benjamini-Hochberg method. Differential abundance analysis was conducted using ALDEx2, which employs a combination of Welch’s t-tests, Wilcoxon rank-sum tests, and effect size estimates to identify differentially abundant taxa between groups.

## Supporting information

Gut3Gel_supplementaryInfo

## Acknowledgments

The authors would like to thank the European Innovation Council through the Accelerator mechanism (HORIZON-EIC-2023-ACCELERATOROPEN-01 ID: 190135075) for partially funding the validation of the technology to be employed as intestinal models. The authors would like to thank Silvia Serra and Rachele Giannecchini, for technical GC-MS assistance. We would also like to thank Abdul Mateen and Lucy Jongen for their contribution in depicting the intrinsic gradient structure of Gut^3^Gel, and to the former for his time in reviewing the manuscript.

## Data availability

The data for this study have been deposited in the European Nucleotide Archive (ENA) at EMBL-EBI under accession number PRJEB85164 (https://www.ebi.ac.uk/ena/browser/view/PRJEB85164).

