## Supplementary material for "Gut^3^Gel: A High Throughput Mucus Model for Culturing Human Intestinal Microbiota": Gut3Gel_supplementaryInfo

Table S1: Pairwise permutational multivariate analysis of variance (PERMANOVA) of beta-diversity across culturing platforms, based on a weighted UniFrac distance matrix.

| **Comparison** | **adjusted p-value** | **Donor** |
|---|---|---|
| BHI Anaerobic vs BHI Aerobic (*) | 0.002 | Donor 1 |
| Donor vs BHI Aerobic (*) | 0.006 | Donor 1 |
| Donor vs BHI Anaerobic (*) | 0.006 | Donor 1 |
| Gut3Gel vs BHI Aerobic (*) | 0.002 | Donor 1 |
| Gut3Gel vs BHI Anaerobic (*) | 0.002 | Donor 1 |
| Gut3Gel vs Donor (*) | 0.006 | Donor 1 |
| BHI Anaerobic vs BHI Aerobic (*) | 0.002 | Donor 2 |
| Donor vs BHI Aerobic (*) | 0.007 | Donor 2 |
| Donor vs BHI Anaerobic (*) | 0.005 | Donor 2 |
| Gut3Gel vs BHI Aerobic (*) | 0.002 | Donor 2 |
| Gut3Gel vs BHI Anaerobic (*) | 0.002 | Donor 2 |
| Gut3Gel vs Donor | 0.101 | Donor 2 |
| BHI Anaerobic vs BHI Aerobic | 0.485 | Donor 3 |
| Donor vs BHI Aerobic (*) | 0.017 | Donor 3 |
| Donor vs BHI Anaerobic (*) | 0.015 | Donor 3 |
| Gut3Gel vs BHI Aerobic (*) | 0.003 | Donor 3 |
| Gut3Gel vs BHI Anaerobic (*) | 0.003 | Donor 3 |
| Gut3Gel vs Donor (*) | 0.004 | Donor 3 |
| BHI Anaerobic vs BHI Aerobic (*) | 0.002 | Donor 4 |
| Donor vs BHI Aerobic (*) | 0.006 | Donor 4 |
| Donor vs BHI Anaerobic (*) | 0.006 | Donor 4 |
| Gut3Gel vs BHI Aerobic (*) | 0.002 | Donor 4 |
| Gut3Gel vs BHI Anaerobic (*) | 0.002 | Donor 4 |
| Gut3Gel vs Donor (*) | 0.006 | Donor 4 |
| BHI Anaerobic vs BHI Aerobic (*) | 0.002 | Donor 5 |
| Donor vs BHI Aerobic (*) | 0.007 | Donor 5 |
| Donor vs BHI Anaerobic (*) | 0.005 | Donor 5 |
| Gut3Gel vs BHI Aerobic (*) | 0.002 | Donor 5 |
| Gut3Gel vs BHI Anaerobic (*) | 0.002 | Donor 5 |
| Gut3Gel vs Donor (*) | 0.007 | Donor 5 |

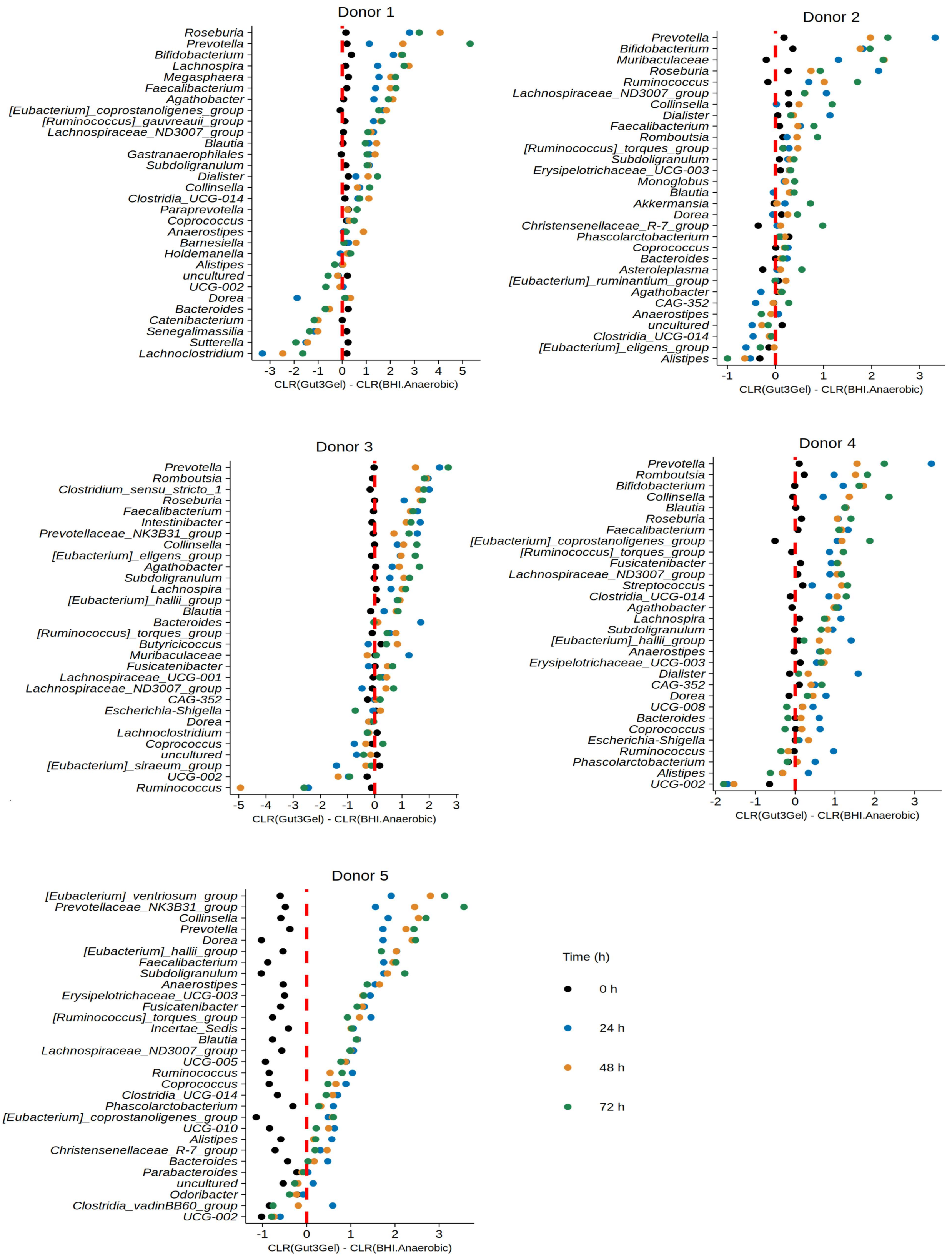


Figure S1: Dotplots displaying the difference between the clr transformed abundances of the Top 30 genera in donor samples in Gut$^3$Gel and BHI anaerobic over the culturing period of 72 hours.

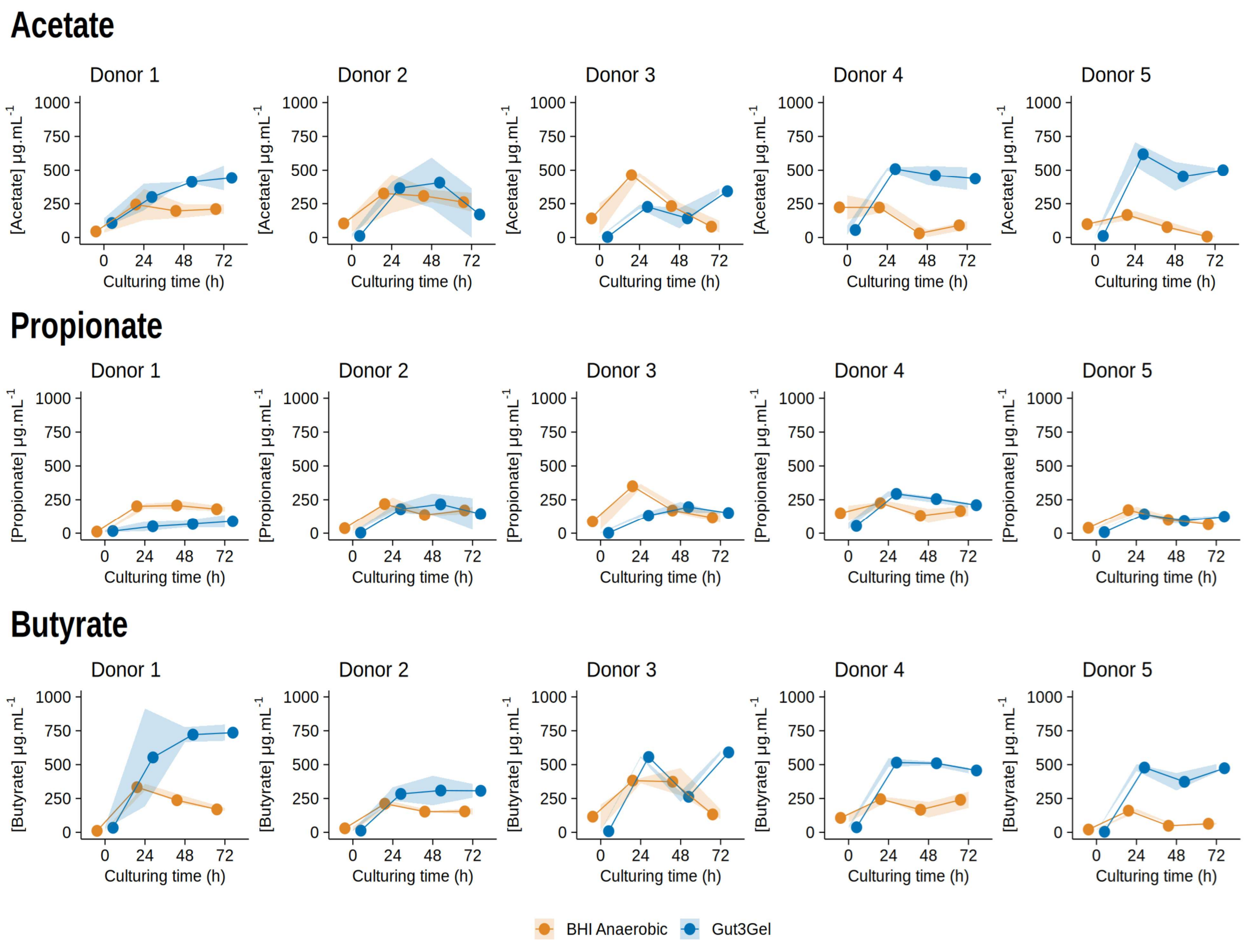


Figure S2: Levels of SCFAs (acetate, propionate and butyrate) produced by the intestinal microbiota inoculated in Gut$^3$Gel, quantified using GC-MS. The data represents the average concentration of each SCFA produced by each donor intestinal microbiota cultured in either Gut$^3$Gel (blue line) or BHI under anaerobic conditions (orange line). The shaded range corresponds to the standard deviation.

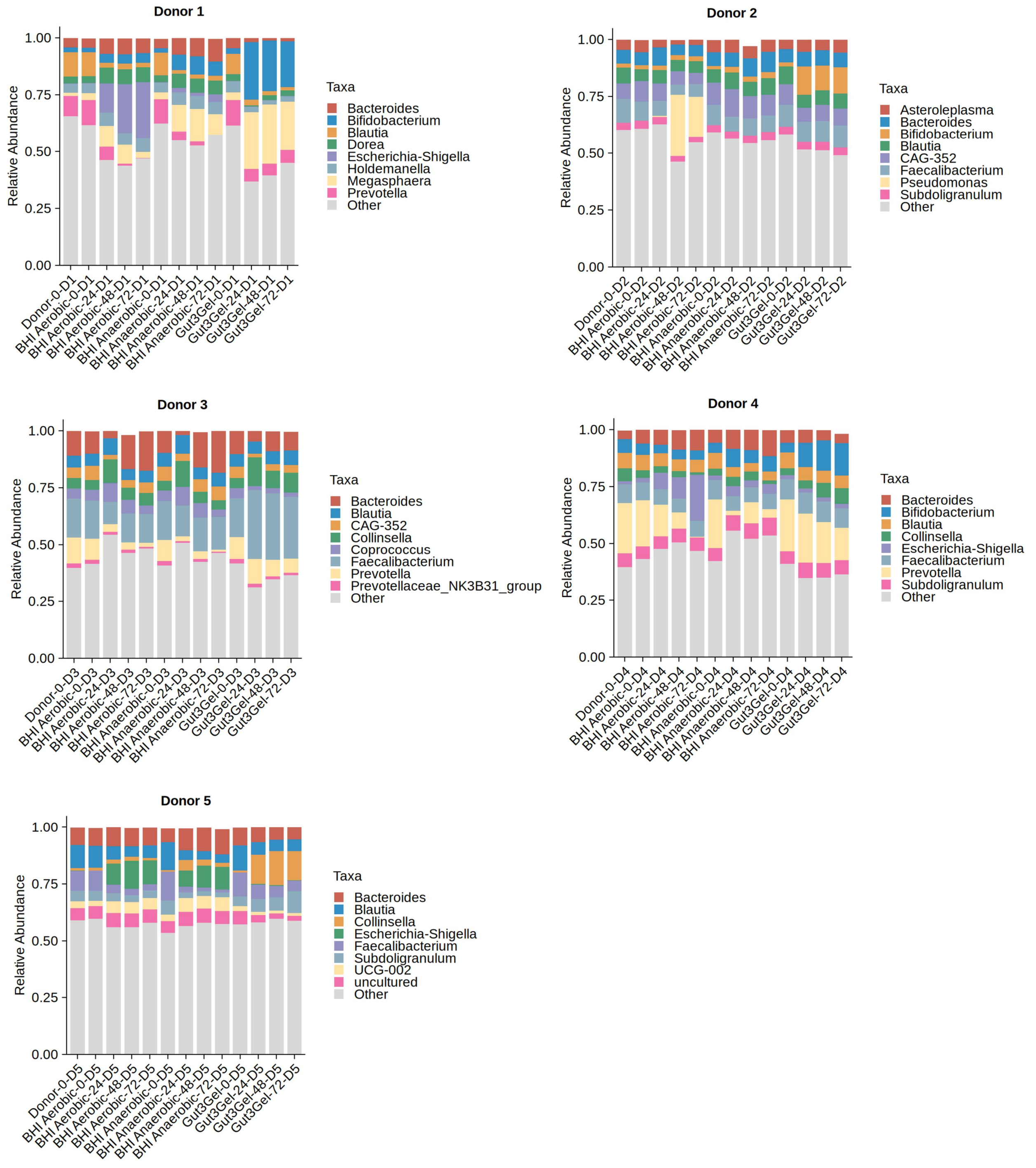


Figure S3: Taxonomic composition at the Genus level across donor samples. The relative abundances are shown for the microbial communities growth in Gut$^3$Gel or BHI (aerobic or anaerobic conditions) cultivated for 24, 48, and 72 hours.
